# Strainy: phasing and assembly of strain haplotypes from long-read metagenome sequencing

**DOI:** 10.1101/2023.01.31.526521

**Authors:** Ekaterina Kazantseva, Ataberk Donmez, Maria Frolova, Mihai Pop, Mikhail Kolmogorov

## Abstract

Bacterial species in microbial communities are often represented by mixtures of strains, distinguished by small variations in their genomes. Despite the functional importance of intra-strain heterogeneity, its characterization from metagenomic sequencing data remains challenging. Short-read approaches can be used to detect small-scale variation between strains, but fail to phase these variants into contiguous haplotypes. Long-read metagenome assemblers can generate contiguous bacterial chromosomes, but often suppress strain-level variation in favor of species-level consensus. Here we present Strainy - an algorithm for strain-level metagenome assembly and phasing from Nanopore and HiFi reads. Strainy takes a de novo metagenomic assembly as input, identifies strain variants which are then phased and assembled into contiguous haplotypes. Using simulated and mock Nanopore and HiFi metagenome data, we show that Strainy assembles accurate and complete strain haplotypes, outperforming current Nanopore-based methods and comparable with HiFi-based algorithms in completeness and accuracy. We then use Strainy to assemble strain haplotypes of a complex environmental metagenome, revealing distinct mutational patterns in bacterial species.

## Introduction

Metagenomic sequencing of human microbiome and other complex microbial communities revealed extensive heterogeneity on sub-species levels (Zhao et al. 2019). Different strains of the same bacterial species often show distinct phenotypes, such as pathogenicity to humans (Kaper, Nataro, and Mobley 2004) or antimicrobial resistance (Schloissnig et al. 2013). Changes in strain composition may provide insights into microbiome evolution (Good et al. 2017), disease associations (Yan et al. 2020) and drug response (Zimmermann et al. 2019).

Strain-level variation could be profiled by mapping of short-read metagenomic data against reference genomes (Albanese and Donati 2017; Olm et al. 2021). However, it is difficult to phase these variants into longer haplotypes due to the limited read length. Alternatively, short-read metagenomic assembly can overcome the reference bias and improve de novo strain detection (Quince et al. 2021; Ghurye et al. 2019), but the assembled genomes are usually fragmented (D. Li et al. 2015; Nurk et al. 2017).

In contrast to short-read methods, long-read metagenomics can recover complete (or nearly-complete) microbial genomes from complex communities (Bertrand et al. 2019; Kim, Ma, and Lee 2022; Dai et al. 2022; Beaulaurier et al. 2020; Van Goethem et al. 2021). Long-read sequencing therefore has better potential to generate contiguous strain haplotypes; however, long-read de novo assemblies typically represent the species-level consensus, suppressing strain heterogeneities. This is because assembly algorithms, such as Canu (Koren et al. 2017), Flye (Kolmogorov et al. 2019) or Shasta (Shafin et al. 2020) were originally designed for noisy long reads with error rates above 10%. Recent studies highlighted that reconstruction of heterogeneous bacterial species (represented by multiple strains) is particularly challenging for long-read assemblers (Kolmogorov et al. 2020; Bickhart et al. 2022; Meyer et al. 2022).

Recently developed HiFi assembly algorithms, such as hifiasm (Cheng et al. 2021), HiCanu (Nurk et al. 2020) and Verkko (Rautiainen et al. 2023) take advantage of highly accurate reads to use (rather than suppress) heterozygous variants to generate diploid human assemblies. This approach was extended in hifiasm-meta (X. Feng et al. 2022) to generate finer metagenome assemblies. Another HiFi-based method called MAGPhase can phase multiple strain haplotypes through the SNP clustering approach (Bickhart et al. 2022). Further, the recently introduced strainFlye pipeline (Fedarko, Kolmogorov, and Pevzner 2022) can detect rare strain mutations in HiFi metagenomic sequencing. HiFi-based methods however are not applicable to the Nanopore sequencing technology due to its higher error rates. Nanopore technology has an advantage of lower cost and better scalability (Kolmogorov et al. 2023); and it was the technology of choice in recent large-scale metagenomic studies (Chen et al. 2022; Jin et al. 2023).

Phasing can be used to complement de novo assembly. In contrast to short reads, long-read sequencing can phase megabase-sized human haplotypes (Martin et al. 2016; Edge, Bafna, and Bansal 2017; Shafin et al. 2021). New algorithms were also developed for polyploid genomes phasing (Schrinner et al. 2020), or assembly graph-based phasing (Chin et al. 2016; Garg et al. 2020; Faure, Guiglielmoni, and Flot 2021). However, these phasing methods were designed for diploid or polyploid genomes and are difficult to apply for metagenomic phasing (Nicholls et al. 2021), since the number of strains and their proportions in a metagenome is not known a priori.

Recently, the authors of the Strainberry algorithm (Vicedomini et al. 2021) introduced an approach for phasing multiple bacterial strains, in which a pairwise phasing algorithm is used iteratively. However, the pairwise approach may not account for uneven distribution of heterozygosity within multiple strains, leading to misassemblies. In another work, the authors of iGDA (Z. Feng et al. 2021) made an improvement in detecting strain SNPs from Nanopore data, but the method does not yield haplotype assemblies. Other studies explored the variant composition of diverse viral populations (Knyazev et al. 2021) using assembly (Jablonski and Beerenwinkel 2021) or long-read phasing (Warwick-Dugdale et al. 2019; Zhou et al. 2018). However, these methods cannot be directly applied to bacterial genomes.

Here we developed Strainy - an algorithm for strain-level metagenome assembly and phasing from both Nanopore and HiFi reads. Using mock, simulated and real Nanopore and HiFi metagenomic data, we show that Strainy assembles accurate and complete strain haplotypes, outperforming current Nanopore-based methods and comparable with the HiFi-based algorithms in completeness and accuracy. We then demonstrate that strain-level assemblies could be used to study distinct patterns of species evolution in a complex activated sludge metagenome sequenced using Nanopore.

## Results

### Overview of the Strainy approach

Strainy could be run either as a standalone phasing approach, or a complete de novo metagenome assembly pipeline called StrainyMAG. The pipeline begins with a de novo assembly (using metaFlye) that produces an assembly graph with collapsed representation of strains. Original reads are then mapped against the assembly graph and the alignment is used to call variable positions. StrainyMAG then performs species-level binning, resulting in metagenome-assembled genomes (MAGs). Then, the Strainy phasing algorithm is run independently for each MAG, generating strain-resolved assemblies of the corresponding MAG species (Figure 1; Methods).

**Figure 1.**
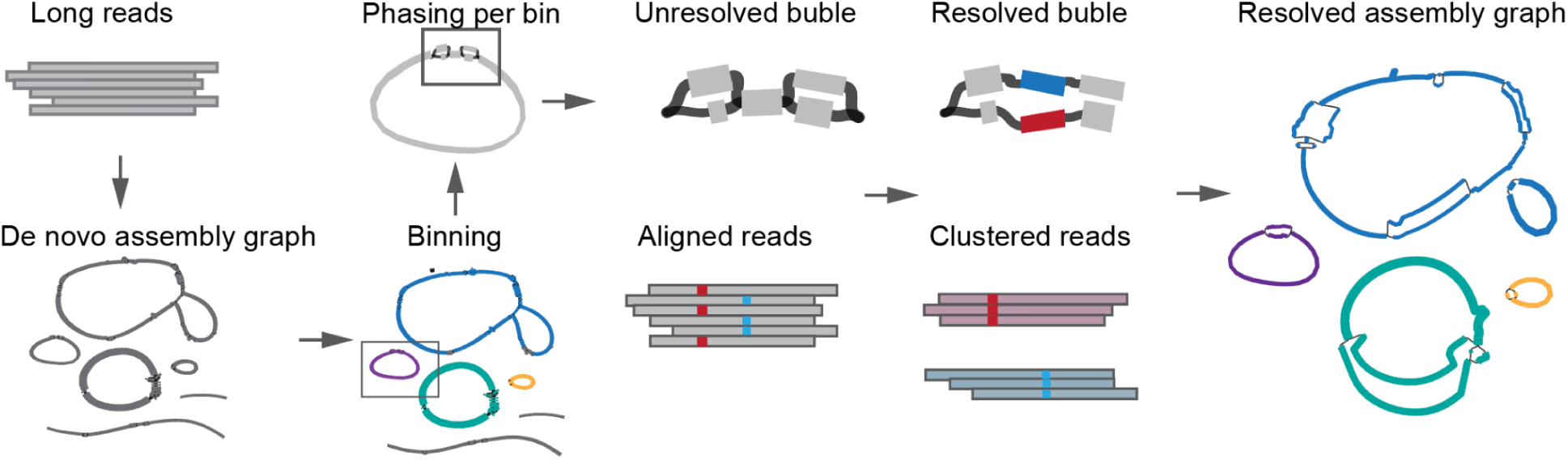
Overview of the StrainyMAG pipeline. Long reads (either Nanopore or HiFi) are de novo assembled and binned into metagenome-assembled genomes (MAGs). The Strainy algorithm is applied to each MAG independently. Strainy aligns reads against the MAG contigs, identifies regions with collapsed strains and phases them into strain-resolved haplotigs. Haplotigs and strain-specific read connections are then used to update and simplify the original de novo assembly graph.

For each contig within a MAG, Strainy builds a *connection* graph, which encodes the pairwise distances between reads aligned to this contig. Next, it clusters reads based on the strain of origin using the community detection approach. While initial clustering separates the most divergent strains, closely related strains (with less variants between them) may remain collapsed. To overcome this, Strainy recursively repeats the clustering procedure with the increased sensitivity to strain variants (Figure 2; Supplementary Figure 1; Methods).

**Figure 2.**
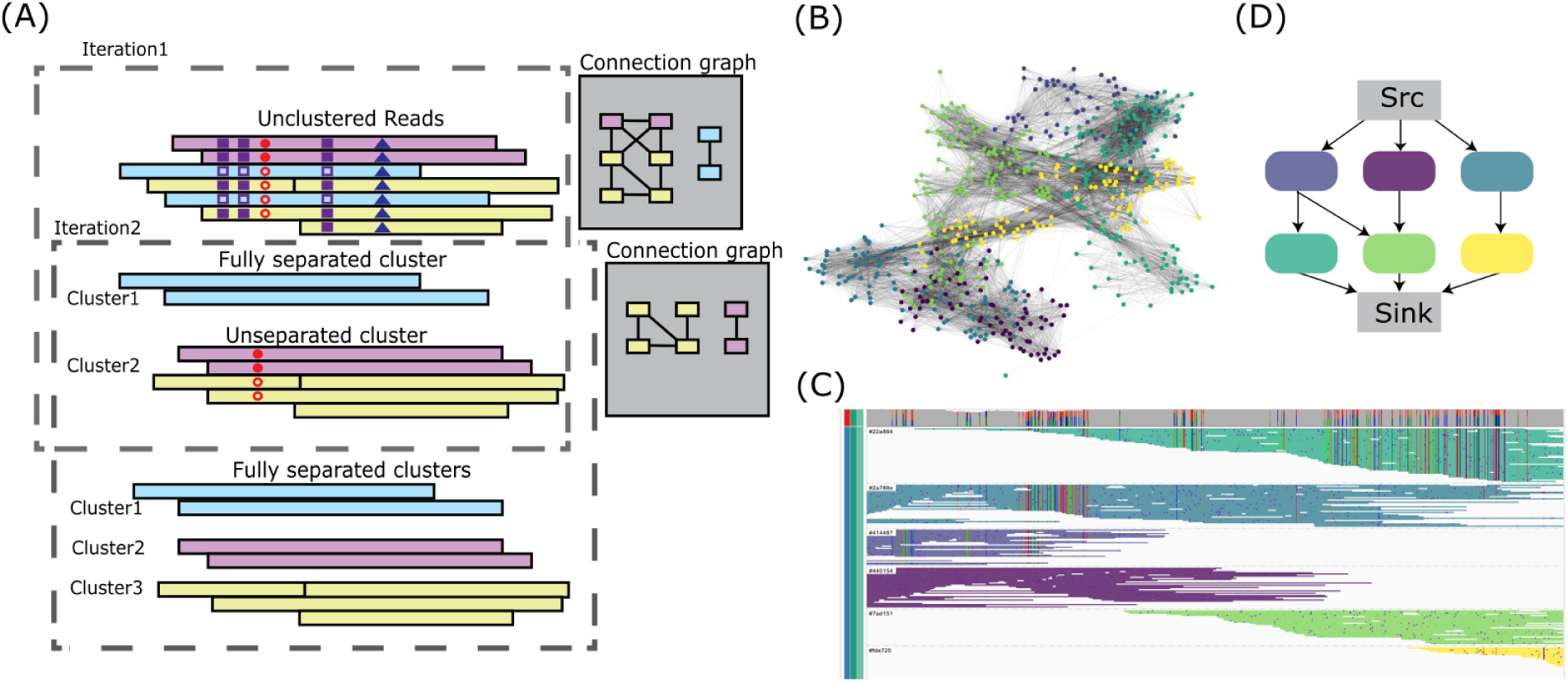
Long-read metagenome phasing algorithm. (A) Outline of the strain clustering algorithm. Given aligned reads, Strainy calls strain-specific SNPs (shown with shapes; genotypes shown with shadings), and builds a connection graph reflecting pairwise read similarity. Strainy then performs clustering via community detection. Clustering procedure is repeated with a reduced set of strain SNPs until convergence. The final clusters represent reads originating from single strains. (B-D) Visualization of strain clustering with real reads from 3 *E.coli* strains. Connection graph clusters (B) separate reads visualized with IGV (C) into phasing groups (shown in different colors). Strainy locally reassembles each strain group, and then builds an overlap graph (D) representing stain haplotigs.

Next, the clustered reads are reassembled locally using the Flye polisher (Kolmogorov et al. 2019), forming strain haplotigs. Strainy then builds an overlap graph of these haplotigs, where paths on the graph correspond to phased strain sequence, interleaved by unphased sequences. Finally, Strainy applies graph simplification algorithms to improve the assembly contiguity.

### Benchmarking using Nanopore and HiFi sequencing of mock bacterial communities

We first used long-read sequencing of mock bacterial communities data to benchmark Strainy along with metaFlye, Strainberry and iGDA. We also benchmarked hifiasm-meta on PacBio HiFi data. We downloaded publicly available metagenomic sequencing of the Zymo mock community D6331, which includes five different *E. coli* strains, along with other bacterial species. We used both HiFi sequencing (https://github.com/PacificBiosciences/pb-metagenomics-tools/blob/master/docs/PacBio-Data.md) and Nanopore sequencing (Liu et al. 2022) for our analysis. Upon inspection of metaFlye and hifiasm-meta assembly graphs, we found that species other than *E. coli* (that were present in single strains) formed separate connected components, representing complete or nearly complete chromosomes. We, thus, focused our benchmarking on five *E. coli* strains (that formed a single tangled connected component), and selected only the reads aligned to the corresponding *E. coli* strain references for our analysis. The coverage of *E. coli* strains in the HiFi dataset was high (100x+), which is rarely achieved in sequencing of real metagenomic samples. We, thus, downsampled HiFi reads to ∼30x read depth of each *E. coli* strain. Overall, the total length of HiFi reads was 700 Mb and read N50 length was 6.1kb; while Nanopore read length was 1.2 Gb with read N50 length ∼5.5kb. Note that while these mock datasets have bacterial strains in relatively equal proportions, below we explore additional metagenomes with uneven species composition.

Strainy takes either a linear reference or a de novo assembly graph as input. A linear reference is not always available for many real metagenomic communities, especially outside of well-studied environments, e.g. human gut (Stewart et al. 2019). Therefore, in our evaluations we used de novo assembly input for Strainy for a truly de novo strain reconstruction. We used metaFlye assemblies for both baseline evaluation and as input for Strainy. We used the “--meta –keep-haplotypes” options to retain the structural variants (SVs) between strains on the assembly graph. Additionally, we modified metaFlye to improve the reconstruction of smaller strain SVs and used the modified version for both baseline and Strainy benchmarks (Methods). We ran hifiasm-meta using the default parameters, and used “p_ctg” sequence output for evaluations (unless noted otherwise). Strainberry and iGDA require a high-quality reference genome as input; we used *E.coli* strain B1109 as reference and default parameters for either HiFi or Nanopore.

Since the strain reference genomes are available, we used metaQUAST (Mikheenko, Saveliev, and Gurevich 2015) to evaluate the accuracy and completeness of strain assemblies. metaQUAST initially determines the strain of origin of each input contig based on the best alignment against all strain references. We used the “--unique-mapping --reuse-combined-alignments” options to ensure a single contig (either strains-specific or strain-collapsed) can only contribute to the completeness of one of the reference strains. In addition to reporting the total number of misassemblies, we estimated the number of strain switch errors (a chimeric connection between two distinct strains) as the number of interspecies translocations reported by metaQUAST.

On both HiFi and Nanopore datasets, Strainy produced strain assemblies with high completeness (95.7% for HiFi and 83.1% for Nanopore), substantially outperforming Strainberry (58.2% for HiFi, 56.8% for Nanopore) and metaFlye (64.7% for HiFi, 29.8% for Nanopore). Further, Strainy had substantially less errors (7 for HiFi and 10 for Nanopore), compared to Strainberry (94 for HiFi and 86 for Nanopore) and metaFlye (22 for HiFi and 19 for Nanopore). Strainy also had similar or better QV scores and comparable NGA50 (Figure 3; Supplementary Table 1). Figure 3 shows an example of Strainy assembly graph transformation, increasing the total length of graph sequence from 7.98 Mb to 27.21 Mb, and the number of nodes from 1,174 to 11,418.

**Figure 3.**
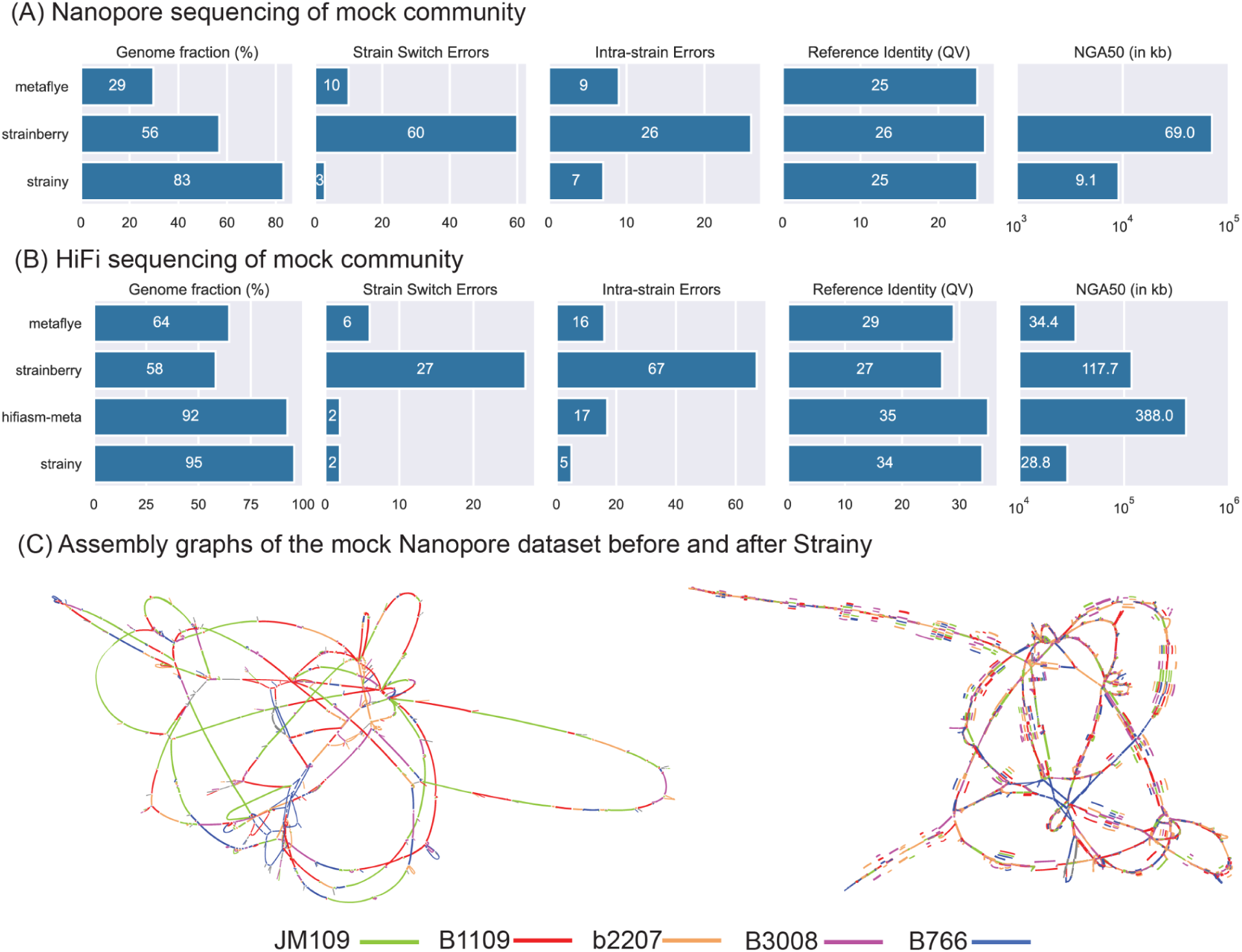
Benchmarking using mock microbial communities. (A, B) Comparison of tools performance using mock communities sequenced with Nanopore R10 and HiFi. Statistics were computed against strain references using metaQUAST with “--unique-mapping” and “--reuse-combined-alignments” options. (C) Assembly graph before and after Strainy transformation (left and right, respectively). After transformation, assembly size increased from 7.98 Mb to 27.21 Mb. Each graph fragment is colored by the strain reference it has the best alignment to.

On HiFi data, hifiasm-meta produced assemblies with slightly lower completeness, as compared to Strainy; but had substantially higher NGA50 (388 kb vs 28.8 kb). This is likely because hifiasm-meta can take advantage of single SNPs to separate haplotypes, while the Strainy approach requires several strain-specific variants. The more aggressive hifiasm-meta approach led to an increased number of misassemblies (19, as compared to 7 for Strainy), and is not applicable to Nanopore data with higher error rates, as compared to HiFi.

Output contigs of iGDA did not show substantial improvement in completeness as compared to the single strain genome for both mock and simulated read assemblies (described below). We therefore excluded iGDA from the further assembly-based analysis.

### Benchmarking using simulated datasets of various species, strain composition and abundance

While mock community analysis is convenient for reference-based benchmarking, it is limited to a single bacterial species and fixed number of strains. To benchmark tools on different bacterial species and strain composition, we generated a range of simulated datasets with different bacterial composition and complexity. We downloaded four sets of complete references from NCBI: five *E. coli* strains (with pairwise ANI ranging from 98.4% to 99.5%), five *S. aureus* strains (ANI ranging from 98.5% to 99.9%), five *L. monocytogenes* (ANI from 98.6% to 99.8%) and five *P. aeruginosa* (ANI from 98.7% to 99.5%). Pairwise ANIs were computed using skani (Shaw and Yu 2023). For each bacterial species, we created a benchmarking set with 2-5 strains, with both uniform read depth (30x) or linearly decreasing “staggered” depth (e.g. 50x, 40x, 30x for a 3-strain dataset). Overall, this resulted in 4 (species) x 4 (strain number) x 2 (depth mode) x 2 (technology) = 64 datasets (Supplementary Table 2).

We used Badread (Wick et al. 2015) to simulate HiFi and Nanopore reads with different error rate distributions. To mimic the sequencing data of the mock datasets analyzed above, we simulated 0.1% error rate and read length N50 = 5 kb for HiFi; and mean error rate of 3%, and N50 = 5kb for Nanopore.

Similar to the mock analysis, we computed completeness and accuracy statistics using metaQUAST. Each dataset consists of 2-5 strains, and below we report statistics for both individual strains and aggregate by datasets (Figure 4; Supplementary Figure 1; Supplementary Table 3). On the Nanopore data, completeness of individual strains was substantially higher for Strainy (mean completeness 87.7%), with Strainberry being the second best (mean completeness 71.5%), while metaFlye assembled substantially less unique sequence (mean completeness 34.4%). Strainy was considerably more accurate, compared to Strainberry (152 and 314 total misassemblies, respectively); metaFlye had a comparable, but lower number of misassemblies (total 34), which is explained by substantially lower length of the metaFlye assemblies. HiFi evaluations showed a similar trend, with Strainy and hifiasm-meta having the highest mean completeness (90.6% and 89.3, respectively) and less misassemblies (55 and 45, respectively), as compared to Strainberry and metaFlye (356 and 72, respectively).

**Figure 4.**
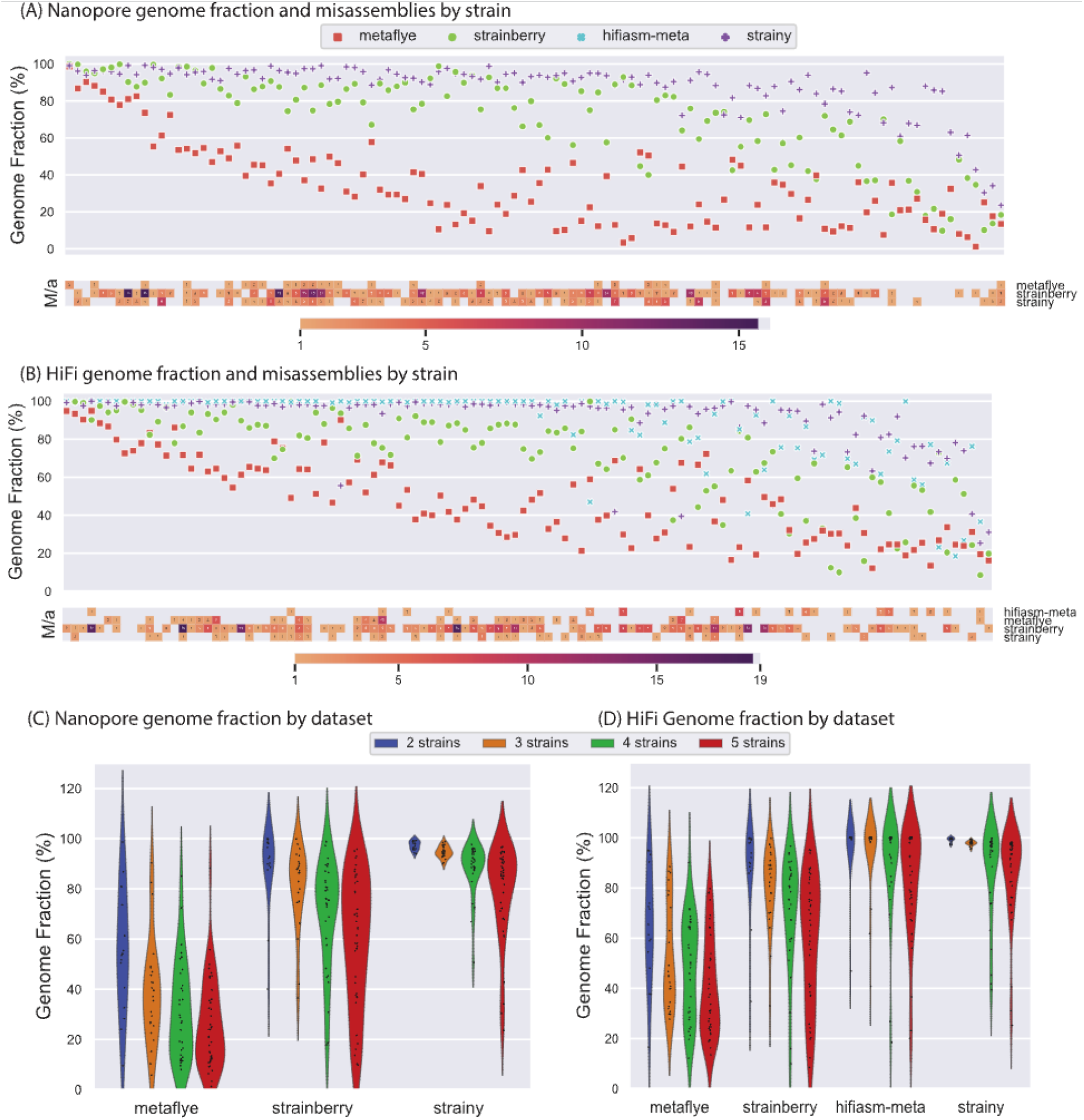
Benchmarking using simulated Nanopore and HiFi sequencing data. (A, B) Assembled strain genome fraction computed using metaQUAST for every bacterial strain in the simulated Nanopore and HiFi datasets. Heatmaps are showing the number of misassemblies for each strain. Strains are sorted in the decreasing mean value among all tools. Corresponding NGA50 plots are available in Supplementary Figure 2. (C, D) Violin plots show distribution of genome fraction aggregated by the number of strains in a simulated dataset.

Aggregated by multi-strain datasets, we observed a trend of decreasing completeness with the increased number of strains for all methods (Figure 4). To explore if any additional features of selected strains affected the performance, we computed pairwise confusion matrices, as the number of inter-strain switches for a given pair of strains. Indeed, strains with higher pairwise average nucleotide identity (ANI) were more likely to have a switch error (Supplementary Figure 3). Interestingly, we did not observe substantial differences between uniform and staggered coverage modes, suggesting that the tools were robust to coverage differences in assembled strains (Supplementary Table 4).

### Assembly and phasing of a complex activated sludge bacterial community with Nanopore and HiFi

We applied Strainy to a metagenomic dataset of activated sludge from an anaerobic digester, sequenced using Nanopore R10 and HiFi sequencing (Sereika et al. 2022). The Nanopore dataset consists of ∼13Gb of reads with N50 ∼6.5 kb. We then applied our StrainyMAG pipeline that first reassembled the data using metaFlye with “--keep-haplotypes” option, resulting in 401Mb assembly and tangled assembly graph with 23,061 nodes. We then realigned reads against the assembly graph unitigs using minimap2 (H. Li 2018), and called SNPs using Clair3 (Zheng et al. 2022). We further applied binning using MetaBAT2 (Kang et al. 2019), which resulted in 32 high-quality MAGs with >80% completeness and <20% contamination. Out of those, 13 MAGs have coverage above the 30x threshold, and were selected for phasing (read depth was varying from 30 to 299).

Overall, phasing of 6 out of 13 MAGs using Strainy resulted in 50-85% of additional sequence for some MAGs, 3 MAGs had moderate increase (10-50%) while 4 MAGs had only a minor increase in sequence (<10%). Those MAGs likely represent single bacterial strains or have alternative strains at very low frequencies. We did not observe strong association between read depth and the amount of sequence increase, suggesting that our MAGs contained one or a few dominant strains, rather than a “continuum” of strains with decreasing abundance. Overall, typically each MAG contained 2-3 strain haplotypes per predicted ORF, but some MAGs had 4-6 distinct strain haplotypes per ORF (Figure 5; Supplementary Table 5). Strain haplotypes (for MAGs with at least two detected strains) had N50 of 8 - 53 kb.

**Figure 5.**
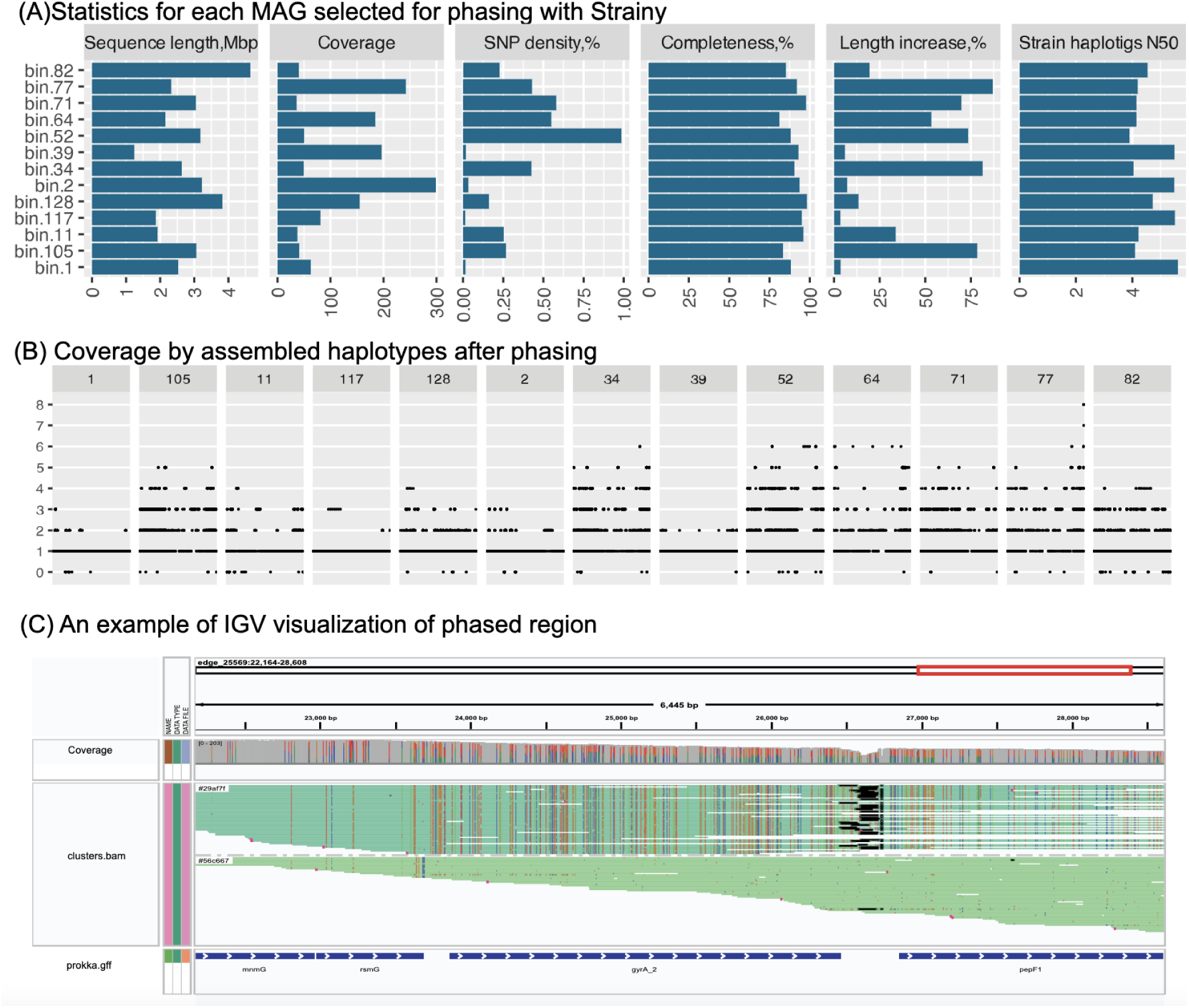
Strain-level assembly of activated sludge metagenome sequenced with Oxford Nanopore R10. (A) Varions statistics for each MAG selected for phasing with Strainy. (B) Coverage of each predicted ORF by assembled Strainy haplotypes. ORFs are ordered according to the coordinates inside unphased MAG sequences. (C) An example of IGV visualization of a phased region of MAG 77 (*Clostridiaceae* family), containing ∼100 bp deletion in the intergenic region and numerous coding SNPs.

Since reference genomes were not available, we instead performed orthogonal validation of assembled strain haplotypes using the HiFi data. We assembled HiFi data using hifiasm-meta, and used the assembled unitigs as input references for QUAST (Supplementary Table 6). As different bacterial species have many homologous regions, we restricted alignment identity to 99% to ensure we are comparing sequences from the similar strains. We also used the unitig (“p_utg”; total size 1.6 Gb), instead of contig output (“p_ctg”; total size 1.2 Gb) of hifiasm-meta, which tends to preserve more strain haplotypes. The length of the aligned sequence (with 99% identity) increased from 32.5 Mb to 46.2 Mb (corresponding to 86% and 81% of the Strainy haplotigs, respectively). The rate of reported misassemblies remained low (2.03 per Mb before phasing; 1.56 per Mb after phasing), confirming high structural concordance between the two technologies.

All MAGs contained indels of size 5-1000 bp, with frequencies following exponential distribution, and primarily placed in intergenic regions (Figure 6). Strain assembly approach - as opposed to short-read variant calling - enables the analysis of phased small and structural variants in conjunction. For example, Figure 5 presents a ∼100 bp deletion in one strain in a non-coding region, flanked by a number of substitutions in surrounding ORFs. Supplementary Figure 4 shows additional examples of complex strain-specific small and structural variants, including examples of SVs altering gene content.

**Figure 6:**
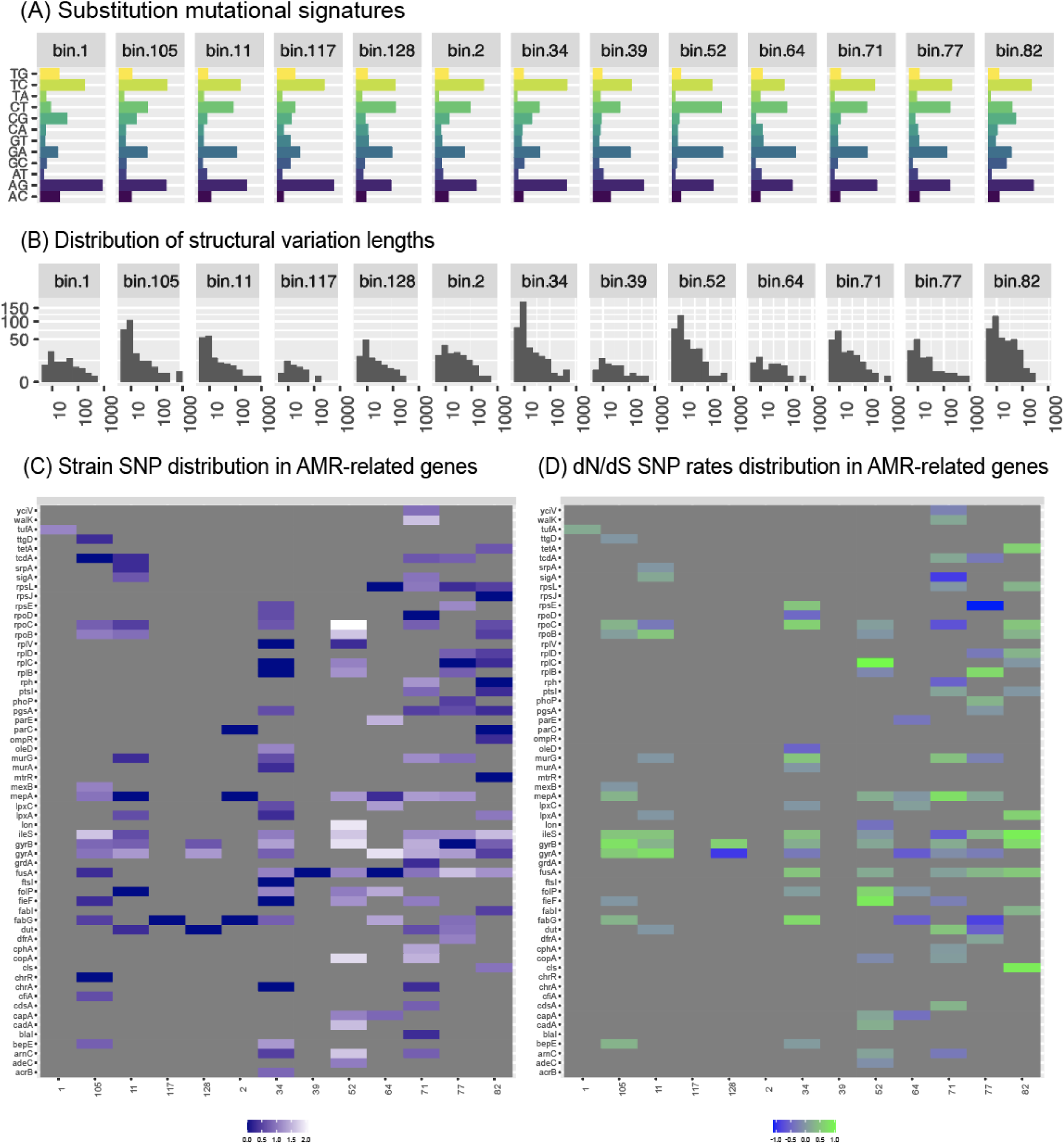
Intra-species small and structural variation provide evolutionary insights. (A) Substitution mutational signatures for each phased MAG (B) Distribution of structural variation lengths for each phased MAG. (C) Frequency (log scale) of substitutions inside AMR-related genes and log-scale ratio of non-synonymous to synonymous mutations.

### Strain haplotypes assemblies provide insights into intra-species evolution

We then proceeded to analyze the strain variants based on the reconstructed haplotypes, to explore species-specific mutational patterns. We aligned Strainy contigs against the MAG consensus as a reference using minimap2, called SNPs and small indels using bcftools (Danecek et al. 2021), and SVs (longer than 50 bp) using hapdiff (Kolmogorov et al. 2023).

The SNP frequency was varying from 0.015% to 0.99% among MAGs (Figure 5). We then performed mutational signature analysis using SNPs, which may highlight different mutational processes (Jee et al. 2016). Substitution signatures were mostly consistent across MAGs (Figure 6), likely reflecting that bacterial cells are likely exposed to the same environment within the metagenome. However, normalized frequencies of some types of substitutions, such as C>G and G>A were variable across MAGs, suggesting species-specific mutational processes.

To explore specific evolution patterns in different species, we focused on the genes from the National Database of Antibiotic Resistant Organisms (NDARO) catalog. A subset of genes with at least one substitution in at least one MAG is shown on Figure 6 (60 out of 9,044 genes in the collection). Each MAG had a characteristic pattern of mutated genes and overall variable mutation rates (Figure 6; Supplementary Table 7). Analysis of non-synonymous to synonymous substitutions rates (dS/dN) revealed a few hotspots with evidence of selection that were specific to MAGs (Figure 6). 32 genes had dS/dN ratio >1; 7 genes had signatures of strong positive selection in more than two MAGs; 2 genes had signatures of both positive and negative (dS/dN < 1) selection in different MAGs (Figure 6).

The selection signatures were consistent with the current literature. For example, mutations in MAG 71 (*Firmicutes*) genes involved in cell wall synthesis are consistent with developing resistance via limiting drug uptake. The mutations in *WalK* contribute to the development of vancomycin resistance in *Staphylococcus aureus* (Zhu et al. 2021), as well as waldiomycin resistance in *Bacillus subtilis* (Kato et al. 2017). *MurG* encodes an enzyme involved in the biosynthesis of peptidoglycan, a crucial component of bacterial cell walls. Positive selection on the *murG* gene was linked to a potential mechanism for the development of vancomycin resistance in *Clostridium difficile* (Leeds et al. 2014).

Active efflux of drugs from bacterial cells is another mechanism of antibiotic resistance. Several MAGs had signatures of positive selection in efflux pump genes *mepA* and *tetA*. Mutations in *mepA* may contribute to multidrug resistance in *Staphylococcus aureus* (Huang et al. 2023), while mutations in *tetA* increase tetracycline resistance in *Escherichia coli* (Jagdmann, Andersson, and Nicoloff 2022).

Most of the NDARO catalog genes with positive selection signatures in our MAGs were housekeeping genes, that despite their essential functions can undergo mutations that confer selective advantages. The high level of quinolone resistance is closely associated with mutations occurring in the *gyrA* in *Bacteroides species* (Oh et al. 2001) and *Escherichia coli* (Shenagari et al. 2018). Mutations in *fusA* have been linked to fusidic acid resistance in *Staphylococcus aureus* (Besier et al. 2003). Mutations in *fabG* can lead to changes in the structure or activity of the β-ketoacyl-ACP reductase enzyme, reducing its susceptibility to triclosan inhibition in *Escherichia coli* (Khan et al. 2016). Mutations in *rpoC* lead to alterations in the RNA polymerase enzyme, affecting multidrug resistance of *Mycobacterium tuberculosis* strains (de Vos et al. 2013). Overall, our analysis highlights how strain-level assemblies can reveal species-specific mutational and selection patterns within a single metagenomic community.

## Discussion

In this work we presented an algorithmic framework for long-read assembly-based phasing of bacterial strains. In contrast to previous approaches, our algorithm can operate on assembly graphs and therefore improve de novo metagenomic assemblies. We implemented the framework into a Strainy phasing tool, and an additional StrainyMAG pipeline for processing of real large metagenomic datasets.

Using mock metagenomic sequencing and simulated data, we showed that Strainy substantially improved over the alternative Nanopore-based approaches for metagenomic strain analysis in terms of completeness and accuracy. Our analysis of simulated datasets with variable strain composition suggests that the presence of closely-related genomes remains one of the main hurdles in long-read metagenomic assembly, in agreement with previous studies (Meyer et al. 2022).

We applied Strainy to assemble strain haplotypes from Nanopore sequencing of an activated sludge bacterial community, and used these haplotypes to analyze intra-species genomic variants. We discovered a substantial number of structural variants in various species, many of which were intergenic, but in some cases resulted in insertions or deletions of one or multiple genes. SNP distribution in AMR genes was characteristic for different species, highlighting the possibility to study bacterial community dynamics and evolution in fine detail.

Strainy substantially outperformed Strainberry in terms of haplotype completeness and accuracy, but was 2-3 times slower (Methods). Strainy running times were nevertheless practical, and we are currently working on an optimized implementation. De novo assemblers (hifiasm-meta and metaFlye) were faster, but did not achieve strain-level resolution of the Nanopore datasets.

On HiFi data, Strainy had comparable completeness, but lower haplotype contiguity, as compared to hifiasm-meta. This is likely because hifiasm-meta is tuned for highly accurate HiFi reads; whereas the Strainy approach is designed to tolerate higher noise of Nanopore reads. In our experiments, we found that imperfections of metaFlye assembly graphs (such as missed adjacencies) often result in breaks in haplotypes. It is likely that improving metaFlye assembly graph concordance and implementing better read-graph alignment will improve the contiguity of Strainy assemblies.

In this work we introduced the haplotype graph framework as a solution for the multi-allele phasing problem, which may not always have linear representation of phased blocks, if heterozygous positions are unevenly distributed in strain genomes. The proposed framework is not limited to bacterial strain phasing, and could be adopted to other applications, for example oncogene amplifications in cancer. We are planning to extend Strainy into a general-purpose multi-allelic phasing framework in our future studies.

## Acknowledgements

This research was supported in part by the Intramural Research Program of the Center for Cancer Research, National Cancer Institute, National Institutes of Health.

## Methods

### Metagenome strain phasing problem

The definition of a bacterial strain could be dependent on the study context (Olm et al. 2021; Bickhart et al. 2022), with some works considering a single nucleotide change in a bacterial genome to be a characteristic of a new strain or lineage. Here, we require strains to have a substantial rate of distinguishing variants, which is a function of the read error profile, read length distribution and sequencing depth. We call bacterial strains that satisfy these requirements for a particular dataset *resolvable*. Our analysis above illustrates that strains with 0.1% heterozygosity and 10x read depth could be resolved with Strainy (with either Nanopore or HiFi sequencing). Although not biologically equivalent, resolvable strains act as haplotypes in the context of metagenomic phasing, and we use these terms interchangeably throughout the paper.

A pairwise phasing problem is defined for a linear reference genome, a set of heterozygous variants and reads that cover contiguous subsets of these variants (Martin et al. 2016; Edge, Bafna, and Bansal 2017). The goal of the phasing procedure is to assign the heterozygous genomic variants to their respective haplotypes. Most phasing algorithms are formulated for two haplotypes, e.g. for a diploid genome. Although some approaches have been extended to polyploid genome phasing, it is assumed that the ploidy is known and haplotypes are represented in equal proportions (Schrinner et al. 2020). Because of that, it is difficult to apply the existing approaches to strain phasing, when the number of strains and their proportions are unknown (Nicholls et al. 2021). Below, we describe an assembly-based phasing algorithm that does not make assumptions about the number of strains or their abundance.

As high-quality references are often not available for uncultured microorganisms, de novo assembled contigs could be used instead. However, long-read assemblers often suppress small differences between strains, and instead generate the species-level consensus. We call contigs representing multiple strains *strain-collapsed*, and contigs that correspond to a single strain *strain-resolved*. Assembly graphs will often contain both strain-resolved and strain-collapsed unitigs which are interleaved, creating additional tangles (for example, bubbles or superbubbles).

Here we use the definition of an assembly graph in which nodes contain assembled sequence, and edges representing adjacencies (and optional dovetail overlaps) between nodes. We call the entire sequence inside a graph node a *unitig*, as opposed to contig that can span multiple assembly graph nodes.

Below we describe the Strainy algorithm, that takes an initial de novo metagenomic assembly as input, and expands the assembly by reconstructing strain haplotypes. In the transformed assembly: (i) each resolvable strain is fully covered by one or multiple contigs, and (ii) each assembled contig corresponds to exactly one resolvable strain. The algorithm consists of two main stages: (i) phase each strain-collapsed unitig into multiple new unitigs that are strain-resolved in a process called *linear phasing* and (ii) simplify and expand the resulting assembly graph.

Strainy phasing algorithm assumes a relatively simple assembly graph with one or a few collapsed bacterial species. For more complex metagenomes, we also provide a StrainyMAG pipeline that performs de novo assembly, and partitions the resulting assembly graph into smaller subgraphs using the MAG partitioning. Strainy is then run on each assembly subgraph independently.

### Informative variants and graph representation of multi-allelic phasing

The goal of the linear phasing stage is to phase each strain-collapsed unitig into a set of strain-resolved unitigs. We denote the input strain-collapsed unitig as *reference* unitig. Let us assume that the reference unitig represents a set of collapsed and resolvable strains *S* of size *|S|*. Each strain *s* ∈ *S* has the abundance of at least *minStrainFreq*. Strain haplotypes are distinguished by their characteristic variants. In this work, we rely on single nucleotide polymorphisms (SNPs) since they typically can be called with much higher confidence (compared to indels) from long reads. A universal set of variants *V* could be computed by calling SNPs with frequencies at least *minStrainFreq* (was set to 0.1 in our benchmarks) from the reference unitig read alignment. Each variant *v* from *V* may be characteristic for one or multiple strains (including all of them in the case of an error in the reference unitig sequence). Therefore, variant *v* splits the set of strains *S* into two subsets, *S_v_* and 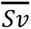, corresponding to strains containing and not containing the variant, respectively.

Given a subset of strains *T* ⊂ *S*, we call variant *v T-informative*, if *v* separates *T* into two non-zero subsets. For example, a variant contained in all strains from *S* is never informative. A variant that is unique to a single strain *s* is always informative for subsets that include *s*. Which variants are informative for each strain subsets is determined by the evolutionary relationship between strains (Supplementary Figure 1). We however do not assume that the strain phylogeny is consistent for all reference unitigs.

Strain variants are often distributed non-uniformly across a genome, creating regions of relatively high and low heterozygosity, with some regions having no variability at all. A long non-heterozygous region is often difficult or impossible to phase. In diploid genomes, phased regions (or haplotype blocks) are interleaved with unphased regions of low heterozygosity. However, in the case of three or more haplotypes a linear definition may be insufficient. Supplementary Figure 1 shows an example of three strains, one of which is more evolutionary distant, compared to two others. While the more distant strain can be fully separated, two other strains share long regions with no heterozygosity.

For example, among the five *E.coli* strains selected for simulations above, we identified 88 regions with no heterozygosity of length longer than 5kb between two *E. coli* strains, while there were 45 such regions shared by three *E. coli* strains and only 3 regions shared by four strains.

### Connection graph and generating strain unitigs

Strainy takes as input long-read alignment using minimap2 (H. Li 2018), which initially does not take assembly graph connectivity into account. Using the bam alignment for each reference unitig, we call SNPs with frequency at least *minStrainFreq* using either mpileup or technology-specific variant caller (such as Clair3).

Next, Strainy clusters long reads based on their strain of origin as follows. Given a set of all reads *R* aligned to a reference unitig (either in full or partially), Strainy builds a *connection* graph. Each node *n* corresponds to a read *r*, and two nodes *n_1_, n_2_*are connected with an undirected edge if the corresponding reads *r_1_* and *r_2_*: (i) have a dovetail overlap of length at least *minOverlap* (typical value = 1000) (ii) overlap spans at least one informative SNP position and (iii) reads have at most *maxSnp* non-matching informative SNP positions (in our benchmarks, *maxSNP* was set to zero for both HiFi and Nanopore). Note that the connection graph construction depends on the informative SNPs, which is a subset of the universal set of SNPs in a reference unitig.

Intuitively, reads from the same strain of origin will form dense connected components within the connection graph (Figure 2). Strainy then partitions the connection graph using the label propagation algorithm (Raghavan, Albert, and Kumara 2007), separating the graph into densely connected clusters.

However, reads from two different strains may also be connected, in case the corresponding overlap spans SNPs that are not informative for these two strains (but are informative for some other strains). This leads to false inter-strain connections and may result in chimeric clusters. Since the number of strains, their evolutionary relationship, and the respective sets of informative SNPs are not known a priori, we use an iterative approach that gradually reduces the set of SNPs to perform fine-grade strain clustering.

At each iteration, for each cluster output by the community detection algorithm, we reduce the set of informative SNPs based only from reads in this cluster. Afterwards, we remove connection graph edges corresponding to overlaps that no longer span any informative SNP positions and repeat the community detection clustering. The process is stopped when the set of informative SNPs does not change substantially. In practice, more than two iterations were rarely needed: on the simulated dataset with five *E. coli* strains sequenced with HiFi reads, ∼80% of clusters required a single iteration, and ∼19% two iterations.

Strainy computes the consensus of each final cluster using the Flye polisher (Kolmogorov et al. 2019), forming strain-resolved unitigs (or *strain* unitigs). Reads from the regions of no heterozygosity will correspond to orphaned nodes in the connection graph. Such reads are separately clustered based only on their reference unitig coordinates. This corresponds to unphased, or *blank* unitigs.

### Strain assembly via overlap graph

Strain unitigs are designed to be highly accurate and represent consensus of reads from a single strain. However, regions of low heterozygosity may reduce density in the connection graph. As a result, the community detection algorithm may split reads from the same strain into multiple strain unitigs (Figure 2).

Strain unitigs that correspond to the adjacent parts of the strain genome will have dovetail overlaps. We then construct an overlap graph of strain unitigs. Two nodes are connected if the corresponding unitigs: (i) overlap by at least *minOverlap* bases; and (ii) their consensus sequences are *sufficiently* similar. The consensus sequences of each node are deemed sufficiently similar if there is no mismatch between the alignment of the sequences and if there are no gaps longer than *maxGap* bases (typical value = 5). The graph also has source and sink nodes, corresponding to the reference unitig endpoints. Strain unitigs originating from reads that align to the start or end of the reference unitig are connected to source or sink nodes, respectively.

In the overlap graph, each strain corresponds to a walk from the source to the sink node. A fully phased strain haplotype corresponds to a non-branching path. However, regions of no heterozygosity between two or more strains will form forks (typically between strain and blank unitigs). Therefore, forks naturally break the fully-phased strain unitigs with unphased regions. The overlap graph therefore defines the final solution to phasing of a reference unitig. Strainy additionally applies the standard simplification procedures, such as: (i) condensing non-branching paths, (ii) removing nested nodes and (iii) transitively reducing bubbles. In addition, short strain unitigs that do not belong to any walk from source to sink are removed. The final graph is denoted as a haplotype graph.

### Assembly graph expansion and simplification

For each (strain-collapsed) reference unitig in the original assembly graph, Strainy performs the linear phasing procedure described above, resulting in a haplotype graph. Afterwards, Strainy substitutes each reference unitig with the corresponding haplotype graph. After this operation, the number of strain-resolved unitigs will increase, however some strain-collapsed unitigs will remain. The goal of the final simplification step is to further extend the strain-resolved unitigs, but retain the connections with strain-collapsed unitigs.

Strainy connects two newly formed strain unitigs if (i) they are connected in one of the overlap graphs or (ii) if there are at least *minLink* reads that span both unitigs. The second condition is aimed at retaining the connectivity of the strain-collapsed nodes of the original assembly graph. In practice, we don’t need to realign reads to test the condition, but instead reuse the information from the connection graph. Afterwards, Strainy performs additional simplification procedures on the updated assembly graph. If a node has two outgoing links, it is forming a *fork*. For every fork, we remove one of the outgoing connections if the resulting transformation does not result in new tips.

### Modification of metaFlye to preserve heterozygous structural variation in the assembly graph

Strainy assumes that all regions that require phasing are free of heterozygous structural variations (SVs) between collapsed strains. Therefore, all such SVs must be preserved on the graph, typically corresponding to bubbles, loops or forks. Flye and metaFlye were originally optimized for consensus assembly (either metagenome or diploid genome), and the algorithm collapses bubbles that correspond to heterozygous SVs. Even though –keep-haplotypes mode was introduced in metaFlye, our analysis of assembly graphs of simulated strain datasets revealed that the graphs may still collapse some SVs because of heuristics that were optimized for consensus assembly.

In this work, we switched off or adjusted some of these heuristics when the –keep-haplotypes mode is enabled. In particular, we introduced a hard alignment gap threshold (corresponding to maximum SV size that could be collapsed), which previously was indirectly controlled through the local alignment score. We have also disabled aggressive duplication filtering during disjointig generation, as disjointigs that represent alternative SV haplotypes may have high similarity and be confused with duplications. These and other changes are now incorporated into a new 2.9.3 Flye release.

### StrainyMAG pipeline

Complex metagenomes often contain many bacterial species, which may form separate connected components on the assembly graph, or remain tangled because of unresolved shared sequence. The goal of StrainyMAG pipeline is to partition the original assembly into smaller parts that could be processed independently.

First, StrainyMAG performs de novo assembly using metaFlye with the “--nano-hq –meta –keep-haplotypes” options. Then, original reads are then mapped back against the graph unitigs using minimap2 (H. Li 2018). Heterozygous SNPs are then called using Clair3 (Zheng et al. 2022). Next, StrainyMAG performs binning using MetaBAT2 (Kang et al. 2019), and bins are evaluated for completion and contamination using CheckM2 (Chklovski et al. 2023). By default, high-quality MAGs with >80% completeness and <20% contamination are selected for further phasing, although this could be adjusted by the user. For each MAG, we also compute heterozygous SNP frequency rate. High-quality MAGs with SNP rate at least 0.1% and coverage at least 30x are selected for the further analysis.

Afterwards, StrainyMAG extracts subgraphs that are defined by the metagenomic binning as follows. Each graph unitig may belong to a single MAG, or not included into any MAGs. For each MAG, we start with its unitigs, and then attempt to incorporate smaller unitigs that often represent alternative SV haplotypes and were not binned because of shorter length. For each pair of the binned unitigs, we search for paths in the assembly graph shorter than 8 nodes that do not contain unitigs that belong to any other MAGs. If such path exists, unitigs and links along this path are added to the MAG subgraph. We then subset the original bam alignment (against the whole assembly graph) to the MAG unitigs. The assembly subgraph and reduced bam alignment are then used as Strainy input.

### Computational performance

Strainy took 160 and 660 minutes to assemble the mock metagenomic datasets, while Strainberry took 237 and 247 minutes, HiFi and Nanopore respectively. Both de novo assemblers were substantially faster: metaFlye took 41 and 60 minutes (HiFi and Nanopore) and hifiasm took 21 minutes (HiFi). All tools were run on a single computational node using 8 threads.

For the simulated datasets, on average, Strainy took 228 and 320 minutes, while Strainberry took 79 and 93 minutes, HiFi and Nanopore respectively. Both de novo assemblers were substantially faster: metaFlye took 14 and 20 minutes (HiFi and Nanopore) and hifiasm-meta took 13 minutes (HiFi).

Overall, Strainy was but was 2-3 times slower as compared to Strainberry, but running times were nevertheless practical, and we are currently working on an optimized implementation. De novo assemblers (hifiasm-meta and metaFlye) were faster, but did not achieve strain-level resolution of the Nanopore datasets.

## Code availability

Strainy is freely available at: https://github.com/katerinakazantseva/stRainy. Complete de novo assembly and phasing pipeline for complex metagenomes are available at: https://github.com/katerinakazantseva/MetagenomeStrainy_ONT_pipeline

## Data availability

Nanopore mock community sequencing: PRJNA804004. HiFi mock community sequencing: https://github.com/PacificBiosciences/pb-metagenomics-tools/blob/master/docs/PacBio-Data.md. Activated sludge sequencing: PRJEB48021. National Database of Antibiotic Resistant Organisms: https://www.ncbi.nlm.nih.gov/pathogens/antimicrobial-resistance/. NDARO catalog: https://www.ncbi.nlm.nih.gov/pathogens/antimicrobial-resistance/.

## Supplementary Tables

**Supplementary Table 1. Evaluations of mock community sequencing datasets.**

**Supplementary Table 2. Information about the simulated datasets.**

**Supplementary Table 3. Evaluations of individual strains assembled from simulated reads.**

**Supplementary Table 4. Evaluations of whole datasets assembled from simulated reads.**

**Supplementary Table 5. Information about the activated sludge MAGs selected for phasing.**

**Supplementary Table 6. Concordance between Nanopore and HiFi assemblies of activated sludge.**

**Supplementary Table 7. Distribution of SNPs in AMR-related genes for different MAGs.**

## Supplementary Figures

**Supplementary Figure 1.**
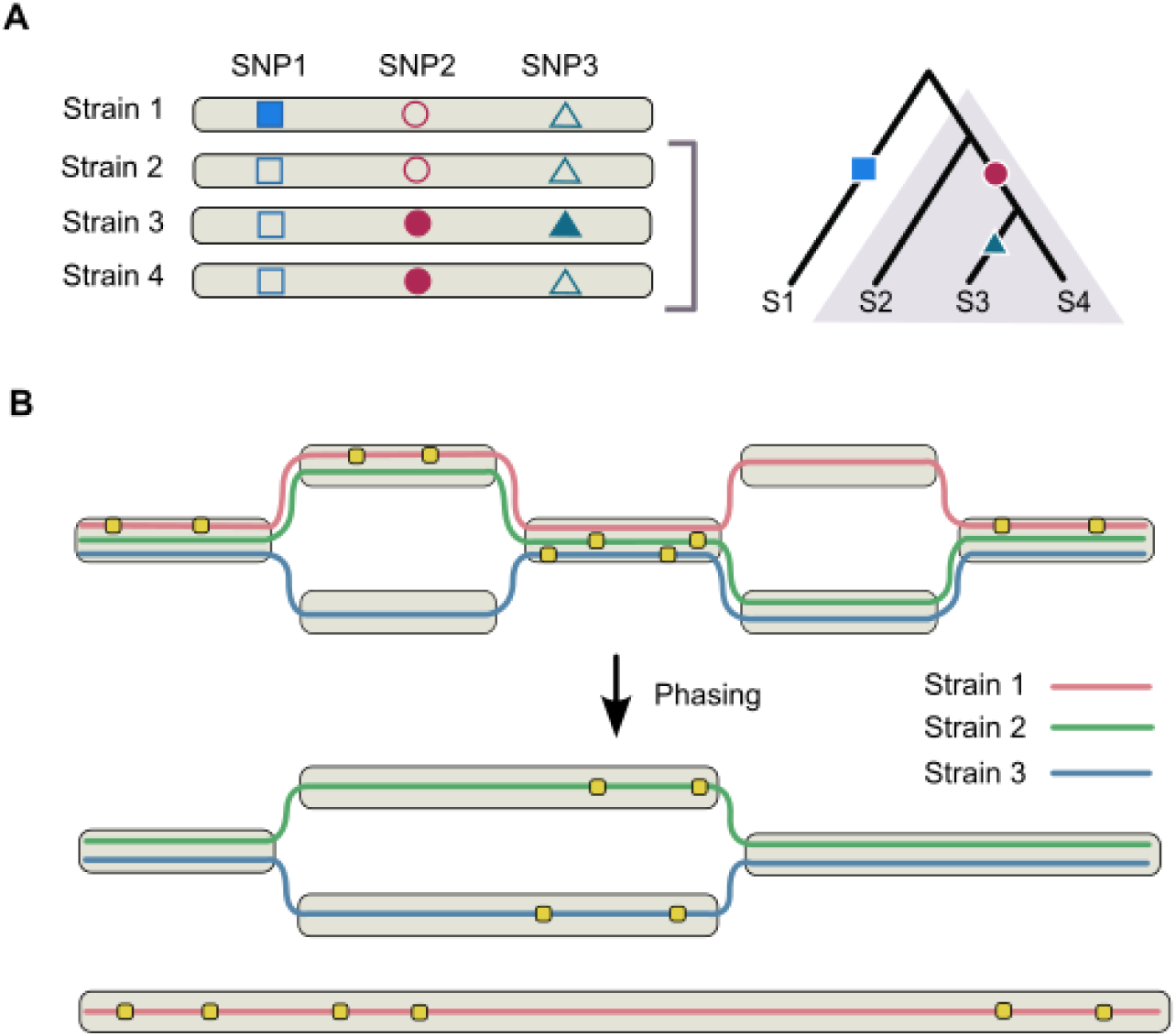
Informative and non-informative SNPs. (A) An example of graph multi-phasing challenge with a set of four closely-related strains and their corresponding phylogenetic tree. SNP positions are shown in different shapes, hollow/solid indicate different genotypes. SNP1 is not informative for the highlighted subset of strains (S2, S3, S4), but SNP2 and SNP3 positions are informative for this subset (B) An example of sequence graph phasing with three strains. Strain-specific variants are shown with yellow rectangles. After phasing and graph simplification, most of the graph nodes are strain-resolved, but some nodes remain strain-collapsed.

**Supplementary Figure 2.**
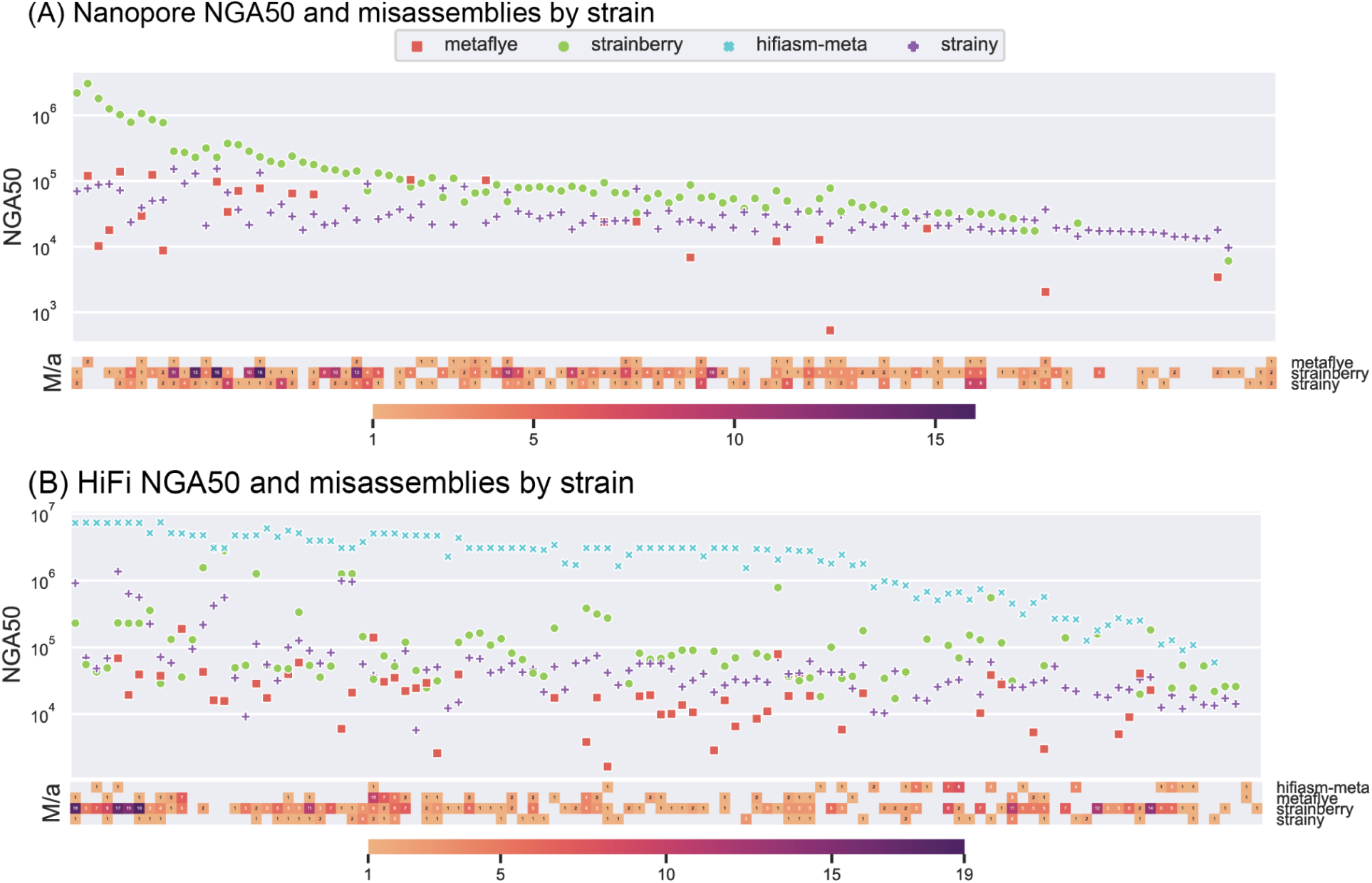
Distribution of NGA50 of the simulated datasets.

**Supplementary Figure 3.**
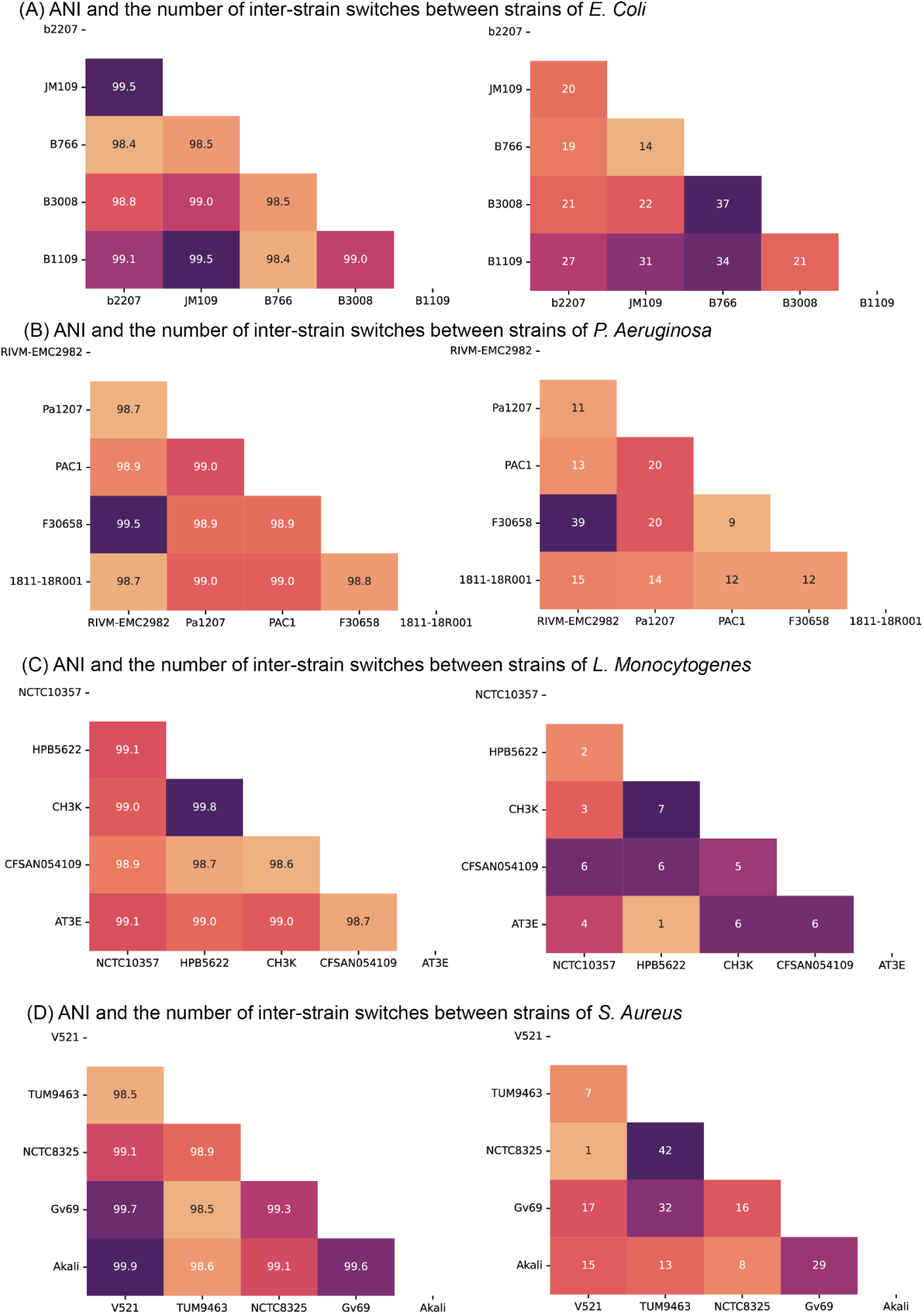
Pairwise average nucleotide identity (ANI) of strain genomes in the simulated datasets and the number of inter-species misassemblies.

**Supplementary Figure 4.**
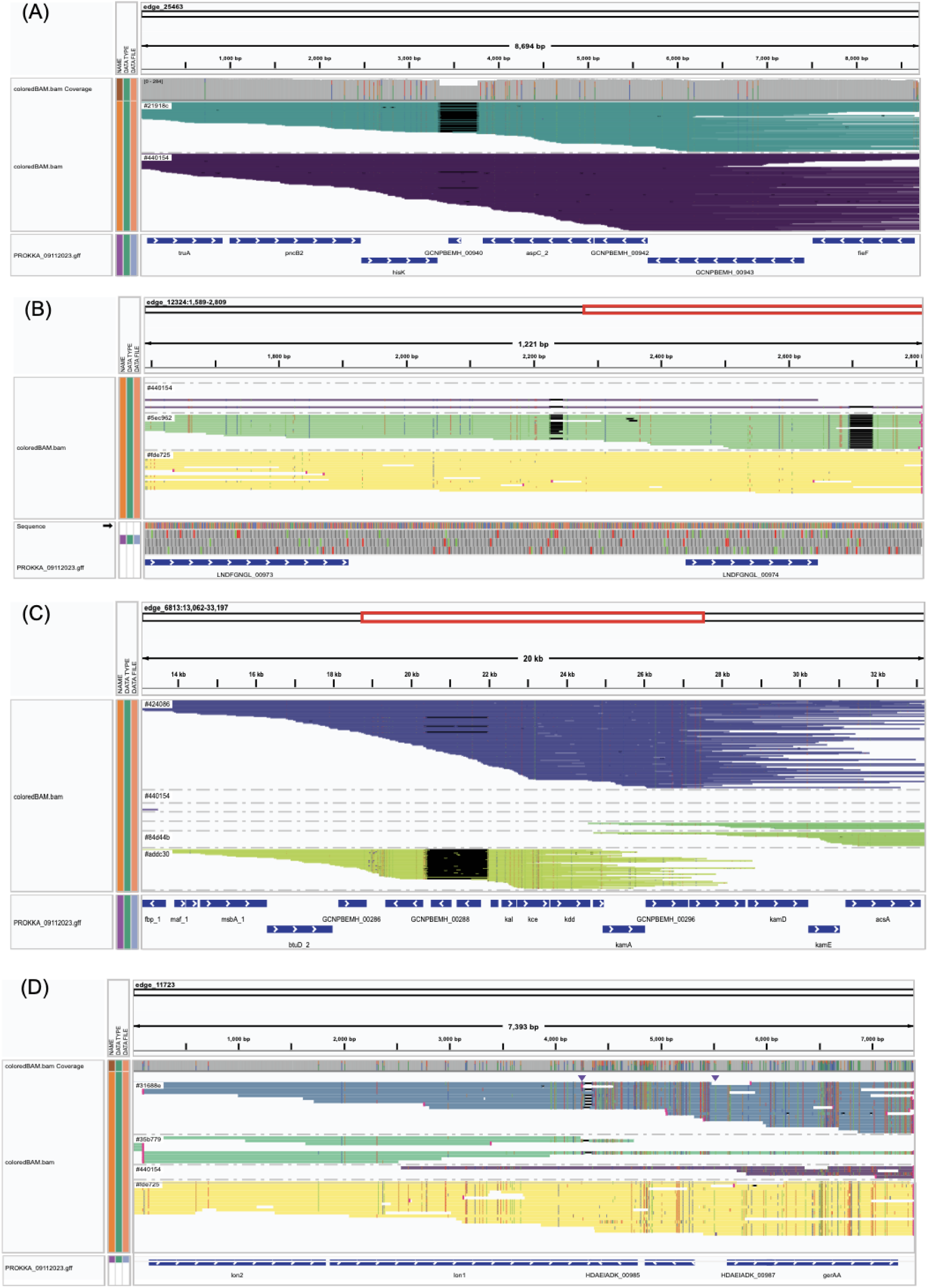
Additional examples of IGV visualization of strain structural variants.

## References

Albanese, Davide, and Claudio Donati. 2017. “Strain Profiling and Epidemiology of Bacterial Species from Metagenomic Sequencing.” Nature Communications 8 (1): 1–14.

Beaulaurier, John, Elaine Luo, John M. Eppley, Paul Den Uyl, Xiaoguang Dai, Andrew Burger, Daniel J. Turner, et al. 2020. “Assembly-Free Single-Molecule Sequencing Recovers Complete Virus Genomes from Natural Microbial Communities.” Genome Research 30 (3): 437–46.

Bertrand, Denis, Jim Shaw, Manesh Kalathiyappan, Amanda Hui Qi Ng, M. Senthil Kumar, Chenhao Li, Mirta Dvornicic, et al. 2019. “Hybrid Metagenomic Assembly Enables High-Resolution Analysis of Resistance Determinants and Mobile Elements in Human Microbiomes.” Nature Biotechnology 37 (8): 937–44.

Besier, Silke, Albrecht Ludwig, Volker Brade, and Thomas A. Wichelhaus. 2003. “Molecular Analysis of Fusidic Acid Resistance in Staphylococcus Aureus.” Molecular Microbiology 47 (2): 463–69.

Bickhart, Derek M., Mikhail Kolmogorov, Elizabeth Tseng, Daniel M. Portik, Anton Korobeynikov, Ivan Tolstoganov, Gherman Uritskiy, et al. 2022. “Generating Lineage-Resolved, Complete Metagenome-Assembled Genomes from Complex Microbial Communities.” Nature Biotechnology 40 (5): 711–19.

Cheng, Haoyu, Gregory T. Concepcion, Xiaowen Feng, Haowen Zhang, and Heng Li. 2021. “Haplotype-Resolved de Novo Assembly Using Phased Assembly Graphs with Hifiasm.” Nature Methods 18 (2): 170–75.

Chen, Liang, Na Zhao, Jiabao Cao, Xiaolin Liu, Jiayue Xu, Yue Ma, Ying Yu, et al. 2022. “Short- and Long-Read Metagenomics Expand Individualized Structural Variations in Gut Microbiomes.” Nature Communications 13 (1): 3175.

Chin, Chen-Shan, Paul Peluso, Fritz J. Sedlazeck, Maria Nattestad, Gregory T. Concepcion, Alicia Clum, Christopher Dunn, et al. 2016. “Phased Diploid Genome Assembly with Single-Molecule Real-Time Sequencing.” Nature Methods 13 (12): 1050–54.

Chklovski, Alex, Donovan H. Parks, Ben J. Woodcroft, and Gene W. Tyson. 2023. “CheckM2: A Rapid, Scalable and Accurate Tool for Assessing Microbial Genome Quality Using Machine Learning.” Nature Methods 20 (8): 1203–12.

Dai, Dongjuan, Connor Brown, Helmut Bürgmann, D. G. Joakim Larsson, Indumathi Nambi, Tong Zhang, Carl-Fredrik Flach, Amy Pruden, and Peter J. Vikesland. 2022. “Long-Read Metagenomic Sequencing Reveals Shifts in Associations of Antibiotic Resistance Genes with Mobile Genetic Elements from Sewage to Activated Sludge.” Microbiome 10 (1): 20.

Danecek, Petr, James K. Bonfield, Jennifer Liddle, John Marshall, Valeriu Ohan, Martin O. Pollard, Andrew Whitwham, et al. 2021. “Twelve Years of SAMtools and BCFtools.” GigaScience 10 (2): giab008.

Edge, Peter, Vineet Bafna, and Vikas Bansal. 2017. “HapCUT2: Robust and Accurate Haplotype Assembly for Diverse Sequencing Technologies.” Genome Research 27 (5): 801–12.

Faure, Roland, Nadège Guiglielmoni, and Jean-François Flot. 2021. “GraphUnzip: Unzipping Assembly Graphs with Long Reads and Hi-C.” bioRxiv. 10.1101/2021.01.29.428779.

Fedarko, Marcus W., Mikhail Kolmogorov, and Pavel A. Pevzner. 2022. “Analyzing Rare Mutations in Metagenomes Assembled Using Long and Accurate Reads.” Genome Research 32 (11-12): 2119–33.

Feng, Xiaowen, Haoyu Cheng, Daniel Portik, and Heng Li. 2022. “Metagenome Assembly of High-Fidelity Long Reads with Hifiasm-Meta.” Nature Methods 19 (6): 671–74.

Feng, Zhixing, Jose C. Clemente, Brandon Wong, and Eric E. Schadt. 2021. “Detecting and Phasing Minor Single-Nucleotide Variants from Long-Read Sequencing Data.” Nature Communications 12 (1): 1–13.

Garg, Shilpa, John Aach, Heng Li, Isaac Sebenius, Richard Durbin, and George Church. 2020. “A Haplotype-Aware de Novo Assembly of Related Individuals Using Pedigree Sequence Graph.” Bioinformatics 36 (8): 2385–92.

Ghurye, Jay, Todd Treangen, Marcus Fedarko, W. Judson Hervey, and Mihai Pop. 2019. “MetaCarvel: Linking Assembly Graph Motifs to Biological Variants.” Genome Biology 20 (1): 1–14.

Good, Benjamin H., Michael J. McDonald, Jeffrey E. Barrick, Richard E. Lenski, and Michael M. Desai. 2017. “The Dynamics of Molecular Evolution over 60,000 Generations.” Nature 551 (7678): 45–50.

Huang, Honghao, Peng Wan, Xinyue Luo, Yixing Lu, Xiaoshen Li, Wenguang Xiong, and Zhenling Zeng. 2023. “Tigecycline Resistance-Associated Mutations in the MepA Efflux Pump in Staphylococcus Aureus.” Microbiology Spectrum 11 (4): e0063423.

Jablonski, Kim Philipp, and Niko Beerenwinkel. 2021. “Computational Methods for Viral Quasispecies Assembly.” In Virus Bioinformatics, 51–64. Boca Raton: Chapman and Hall/CRC.

Jagdmann, Jennifer, Dan I. Andersson, and Hervé Nicoloff. 2022. “Low Levels of Tetracyclines Select for a Mutation That Prevents the Evolution of High-Level Resistance to Tigecycline.” PLoS Biology 20 (9): e3001808.

Jee, Justin, Aviram Rasouly, Ilya Shamovsky, Yonatan Akivis, Susan R. Steinman, Bud Mishra, and Evgeny Nudler. 2016. “Rates and Mechanisms of Bacterial Mutagenesis from Maximum-Depth Sequencing.” Nature 534 (7609): 693–96.

Jin, Hao, Keyu Quan, Qiuwen He, Lai-Yu Kwok, Teng Ma, Yalin Li, Feiyan Zhao, Lijun You, Heping Zhang, and Zhihong Sun. 2023. “A High-Quality Genome Compendium of the Human Gut Microbiome of Inner Mongolians.” Nature Microbiology 8 (1): 150–61.

Kang, Dongwan D., Feng Li, Edward Kirton, Ashleigh Thomas, Rob Egan, Hong An, and Zhong Wang. 2019. “MetaBAT 2: An Adaptive Binning Algorithm for Robust and Efficient Genome Reconstruction from Metagenome Assemblies.” PeerJ 7 (July): e7359.

Kaper, James B., James P. Nataro, and Harry L. Mobley. 2004. “Pathogenic Escherichia Coli.” Nature Reviews. Microbiology 2 (2): 123–40.

Kato, Akinori, Shuhei Ueda, Taku Oshima, Yoichi Inukai, Toshihide Okajima, Masayuki Igarashi, Yoko Eguchi, and Ryutaro Utsumi. 2017. “Characterization of H-Box Region Mutants of WalK Inert to the Action of Waldiomycin in Bacillus Subtilis.” The Journal of General and Applied Microbiology 63 (4): 212–21.

Khan, Raees, Hyun Gi Kong, Yong-Hoon Jung, Jinhee Choi, Kwang-Yeol Baek, Eul Chul Hwang, and Seon-Woo Lee. 2016. “Triclosan Resistome from Metagenome Reveals Diverse Enoyl Acyl Carrier Protein Reductases and Selective Enrichment of Triclosan Resistance Genes.” Scientific Reports 6 (August): 32322.

Kim, Chan Yeong, Junyeong Ma, and Insuk Lee. 2022. “HiFi Metagenomic Sequencing Enables Assembly of Accurate and Complete Genomes from Human Gut Microbiota.” Nature Communications 13 (1): 6367.

Knyazev, Sergey, Lauren Hughes, Pavel Skums, and Alexander Zelikovsky. 2021. “Epidemiological Data Analysis of Viral Quasispecies in the next-Generation Sequencing Era.” Briefings in Bioinformatics 22 (1): 96–108.

Kolmogorov, Mikhail, Derek M. Bickhart, Bahar Behsaz, Alexey Gurevich, Mikhail Rayko, Sung Bong Shin, Kristen Kuhn, et al. 2020. “metaFlye: Scalable Long-Read Metagenome Assembly Using Repeat Graphs.” Nature Methods 17 (11): 1103–10.

Kolmogorov, Mikhail, Kimberley J. Billingsley, Mira Mastoras, Melissa Meredith, Jean Monlong, Ryan Lorig-Roach, Mobin Asri, et al. 2023. “Scalable Nanopore Sequencing of Human Genomes Provides a Comprehensive View of Haplotype-Resolved Variation and Methylation.” Nature Methods 20 (10): 1483–92.

Kolmogorov, Mikhail, Jeffrey Yuan, Yu Lin, and Pavel A. Pevzner. 2019. “Assembly of Long, Error-Prone Reads Using Repeat Graphs.” Nature Biotechnology 37 (5): 540–46.

Koren, Sergey, Brian P. Walenz, Konstantin Berlin, Jason R. Miller, Nicholas H. Bergman, and Adam M. Phillippy. 2017. “Canu: Scalable and Accurate Long-Read Assembly via Adaptive K-Mer Weighting and Repeat Separation.” Genome Research 27 (5): 722–36.

Leeds, Jennifer A., Meena Sachdeva, Steve Mullin, S. Whitney Barnes, and Alexey Ruzin. 2014. “In Vitro Selection, via Serial Passage, of Clostridium Difficile Mutants with Reduced Susceptibility to Fidaxomicin or Vancomycin.” The Journal of Antimicrobial Chemotherapy 69 (1): 41–44.

Li, Dinghua, Chi-Man Liu, Ruibang Luo, Kunihiko Sadakane, and Tak-Wah Lam. 2015. “MEGAHIT: An Ultra-Fast Single-Node Solution for Large and Complex Metagenomics Assembly via Succinct de Bruijn Graph.” Bioinformatics 31 (10): 1674–76.

Li, Heng. 2018. “Minimap2: Pairwise Alignment for Nucleotide Sequences.” Bioinformatics 34 (18): 3094–3100.

Liu, Lei, Yu Yang, Yu Deng, and Tong Zhang. 2022. “Nanopore Long-Read-Only Metagenomics Enables Complete and High-Quality Genome Reconstruction from Mock and Complex Metagenomes.” Microbiome 10 (1): 1–7.

Martin, Marcel, Murray Patterson, Shilpa Garg, Sarah O. Fischer, Nadia Pisanti, Gunnar W. Klau, Alexander Schöenhuth, and Tobias Marschall. 2016. “WhatsHap: Fast and Accurate Read-Based Phasing.” bioRxiv. 10.1101/085050.

Meyer, Fernando, Adrian Fritz, Zhi-Luo Deng, David Koslicki, Till Robin Lesker, Alexey Gurevich, Gary Robertson, et al. 2022. “Critical Assessment of Metagenome Interpretation: The Second Round of Challenges.” Nature Methods 19 (4): 429–40.

Mikheenko, Alla, Vladislav Saveliev, and Alexey Gurevich. 2015. “MetaQUAST: Evaluation of Metagenome Assemblies.” Bioinformatics 32 (7): 1088–90.

Nicholls, Samuel M., Wayne Aubrey, Kurt De Grave, Leander Schietgat, Christopher J. Creevey, and Amanda Clare. 2021. “On the Complexity of Haplotyping a Microbial Community.” Bioinformatics 37 (10): 1360–66.

Nurk, Sergey, Dmitry Meleshko, Anton Korobeynikov, and Pavel A. Pevzner. 2017. “metaSPAdes: A New Versatile Metagenomic Assembler.” Genome Research 27 (5): 824–34.

Nurk, Sergey, Brian P. Walenz, Arang Rhie, Mitchell R. Vollger, Glennis A. Logsdon, Robert Grothe, Karen H. Miga, Evan E. Eichler, Adam M. Phillippy, and Sergey Koren. 2020. “HiCanu: Accurate Assembly of Segmental Duplications, Satellites, and Allelic Variants from High-Fidelity Long Reads.” Genome Research 30 (9): 1291–1305.

Oh, H., N. El Amin, T. Davies, P. C. Appelbaum, and C. Edlund. 2001. “gyrA Mutations Associated with Quinolone Resistance in Bacteroides Fragilis Group Strains.” Antimicrobial Agents and Chemotherapy 45 (7): 1977–81.

Olm, Matthew R., Alexander Crits-Christoph, Keith Bouma-Gregson, Brian A. Firek, Michael J. Morowitz, and Jillian F. Banfield. 2021. “inStrain Profiles Population Microdiversity from Metagenomic Data and Sensitively Detects Shared Microbial Strains.” Nature Biotechnology 39 (6): 727–36.

Quince, Christopher, Sergey Nurk, Sebastien Raguideau, Robert James, Orkun S. Soyer, J. Kimberly Summers, Antoine Limasset, A. Murat Eren, Rayan Chikhi, and Aaron E. Darling. 2021. “STRONG: Metagenomics Strain Resolution on Assembly Graphs.” Genome Biology 22 (1): 1–34.

Raghavan, Usha Nandini, Réka Albert, and Soundar Kumara. 2007. “Near Linear Time Algorithm to Detect Community Structures in Large-Scale Networks.” Physical Review. E, Statistical, Nonlinear, and Soft Matter Physics 76 (3 Pt 2): 036106.

Rautiainen, Mikko, Sergey Nurk, Brian P. Walenz, Glennis A. Logsdon, David Porubsky, Arang Rhie, Evan E. Eichler, Adam M. Phillippy, and Sergey Koren. 2023. “Telomere-to-Telomere Assembly of Diploid Chromosomes with Verkko.” Nature Biotechnology 41 (10): 1474–82.

Schloissnig, Siegfried, Manimozhiyan Arumugam, Shinichi Sunagawa, Makedonka Mitreva, Julien Tap, Ana Zhu, Alison Waller, et al. 2013. “Genomic Variation Landscape of the Human Gut Microbiome.” Nature 493 (7430): 45–50.

Schrinner, Sven D., Rebecca Serra Mari, Jana Ebler, Mikko Rautiainen, Lancelot Seillier, Julia J. Reimer, Björn Usadel, Tobias Marschall, and Gunnar W. Klau. 2020. “Haplotype Threading: Accurate Polyploid Phasing from Long Reads.” Genome Biology 21 (1): 1–22.

Sereika, Mantas, Rasmus Hansen Kirkegaard, Søren Michael Karst, Thomas Yssing Michaelsen, Emil Aarre Sørensen, Rasmus Dam Wollenberg, and Mads Albertsen. 2022. “Oxford Nanopore R10.4 Long-Read Sequencing Enables the Generation of near-Finished Bacterial Genomes from Pure Cultures and Metagenomes without Short-Read or Reference Polishing.” Nature Methods 19 (7): 823–26.

Shafin, Kishwar, Trevor Pesout, Pi-Chuan Chang, Maria Nattestad, Alexey Kolesnikov, Sidharth Goel, Gunjan Baid, et al. 2021. “Haplotype-Aware Variant Calling with PEPPER-Margin-DeepVariant Enables High Accuracy in Nanopore Long-Reads.” Nature Methods 18 (11): 1322–32.

Shafin, Kishwar, Trevor Pesout, Ryan Lorig-Roach, Marina Haukness, Hugh E. Olsen, Colleen Bosworth, Joel Armstrong, et al. 2020. “Nanopore Sequencing and the Shasta Toolkit Enable Efficient de Novo Assembly of Eleven Human Genomes.” Nature Biotechnology 38 (9): 1044–53.

Shaw, Jim, and Yun William Yu. 2023. “Fast and Robust Metagenomic Sequence Comparison through Sparse Chaining with Skani.” Nature Methods, September, 1–5.

Shenagari, Mohammad, Masoud Bakhtiari, Ali Mojtahedi, and Zahra Atrkar Roushan. 2018. “High Frequency of Mutations in gyrA Gene Associated with Quinolones Resistance in Uropathogenic Escherichia Coli Isolates from the North of Iran.” Iranian Journal of Basic Medical Sciences 21 (12): 1226–31.

Stewart, Robert D., Marc D. Auffret, Amanda Warr, Alan W. Walker, Rainer Roehe, and Mick Watson. 2019. “Compendium of 4,941 Rumen Metagenome-Assembled Genomes for Rumen Microbiome Biology and Enzyme Discovery.” Nature Biotechnology 37 (8): 953–61.

Van Goethem, Marc W., Andrew R. Osborn, Benjamin P. Bowen, Peter F. Andeer, Tami L. Swenson, Alicia Clum, Robert Riley, et al. 2021. “Long-Read Metagenomics of Soil Communities Reveals Phylum-Specific Secondary Metabolite Dynamics.” Communications Biology 4 (1): 1–10.

Vicedomini, Riccardo, Christopher Quince, Aaron E. Darling, and Rayan Chikhi. 2021. “Strainberry: Automated Strain Separation in Low-Complexity Metagenomes Using Long Reads.” Nature Communications 12 (1): 1–14.

Vos, M. de, B. Müller, S. Borrell, P. A. Black, P. D. van Helden, R. M. Warren, S. Gagneux, and T. C. Victor. 2013. “Putative Compensatory Mutations in the rpoC Gene of Rifampin-Resistant Mycobacterium Tuberculosis Are Associated with Ongoing Transmission.” Antimicrobial Agents and Chemotherapy 57 (2): 827–32.

Warwick-Dugdale, Joanna, Natalie Solonenko, Karen Moore, Lauren Chittick, Ann C. Gregory, Michael J. Allen, Matthew B. Sullivan, and Ben Temperton. 2019. “Long-Read Viral Metagenomics Captures Abundant and Microdiverse Viral Populations and Their Niche-Defining Genomic Islands.” PeerJ 7 (April): e6800.

Wick, Ryan R., Mark B. Schultz, Justin Zobel, and Kathryn E. Holt. 2015. “Bandage: Interactive Visualization of de Novo Genome Assemblies.” Bioinformatics 31 (20): 3350–52.

Yan, Yan, Long H. Nguyen, Eric A. Franzosa, and Curtis Huttenhower. 2020. “Strain-Level Epidemiology of Microbial Communities and the Human Microbiome.” Genome Medicine 12 (1): 71.

Zhao, Shijie, Tami D. Lieberman, Mathilde Poyet, Kathryn M. Kauffman, Sean M. Gibbons, Mathieu Groussin, Ramnik J. Xavier, and Eric J. Alm. 2019. “Adaptive Evolution within Gut Microbiomes of Healthy People.” Cell Host & Microbe 25 (5): 656–67.e8.

Zheng, Zhenxian, Shumin Li, Junhao Su, Amy Wing-Sze Leung, Tak-Wah Lam, and Ruibang Luo. 2022. “Symphonizing Pileup and Full-Alignment for Deep Learning-Based Long-Read Variant Calling.” Nature Computational Science 2 (12): 797–803.

Zhou, Zhemin, Nina Luhmann, Nabil-Fareed Alikhan, Christopher Quince, and Mark Achtman. 2018. “Accurate Reconstruction of Microbial Strains from Metagenomic Sequencing Using Representative Reference Genomes.” In Research in Computational Molecular Biology, 225–40. Springer International Publishing.

Zhu, Jiade, Banghui Liu, Xueqin Shu, and Baolin Sun. 2021. “A Novel Mutation of walK Confers Vancomycin-Intermediate Resistance in Methicillin-Susceptible Staphylococcus Aureus.” International Journal of Medical Microbiology: IJMM 311 (2): 151473.

Zimmermann, Michael, Maria Zimmermann-Kogadeeva, Rebekka Wegmann, and Andrew L. Goodman. 2019. “Mapping Human Microbiome Drug Metabolism by Gut Bacteria and Their Genes.” Nature 570 (7762): 462–67.

